# Introgression patterns between house mouse subspecies and species reveal genomic windows of frequent exchange

**DOI:** 10.1101/168328

**Authors:** Kristian Karsten Ullrich, Miriam Linnenbrink, Diethard Tautz

## Abstract

Based on whole genome sequencing data, we have studied the patterns of introgression in a phylogenetically well defined set of populations, sub-species and species of mice (*Mus m. domesticus*, *Mus m. musculus*, *Mus m. castaneus* and *Mus spretus*). We find that many discrete genomic regions are subject to repeated and mutual introgression and exchange. The majority of these regions code for genes that are involved in parasite defense or genomic conflict. They include genes involved in adaptive immunity, such as the MHC region or antibody coding regions, but also genes involved in innate immune reactions of the epidermis. We find also clusters of KRAB zinc finger proteins that control the spread of transposable elements and genes that are involved in meiotic drive. These findings suggest that even well separated populations and species maintain the capacity to exchange genetic material in a special set of evolutionary active genes.

## Introduction

Patterns of introgression are frequently observed between taxa that can still hybridize (Green et al., 2010; Salazar et al., 2010; Song et al., 2011; Staubach et al., 2012; Hedrick, 2013; Liu et al., 2015; Zhang et al., 2016; Stukenbrock, 2016; Kumar et al., 2017, and refs therein). The house mouse (*Mus musculus*) forms a species complex with several described and not yet fully described sub-species (Guénet and Bonhomme, 2003; Phifer-Rixey and Nachman, 2015; Hardouin et al., 2015) that are distributed in allopatric patterns across the whole world but can still hybridize, in particular at secondary contact zones (Sage et al., 1986; Teeter et al., 2008; Janoušek et al., 2012; Turner and Harr, 2014). The sister species *Mus spretus* lives in sympatry with *M. m. domesticus* in Western Europe and can form hybrids with this subspecies, but maintains its own species status. Patterns of introgression between the Western and Eastern house mouse (M. m. *domesticus* and *M. m. musculus*) have been intensively studied along the hybrid zone in Europe (Teeter et al., 2008; Wang et al., 2011; Janoušek et al., 2015). Genome-wide comparisons based on SNP array data have further revealed that introgression of haplotypes can also occur across large distances, possibly mediated through human transport of mice (Staubach et al., 2012). Since these introgressed haplotypes are much larger than the average linkage disequilibrium (LD) blocks in wild populations (Laurie et al., 2007), their spread in distant populations is apparently driven by positive selection, since they would rapidly break up under drift conditions (Staubach et al., 2012). A prominent example of an adaptive introgression of a *M. spretus* related haplotype into *M. m. musculus* populations is the locus that confers resistance to the rodenticide Warfarin, *Vkorc1*(Song et al., 2011; Liu et al., 2015). But haplotypes may also introgress between separated populations of the same subspecies, as it has been shown for the MLV virus receptor *Xpr1*(Hasenkamp et al., 2015).

A variety of tests have been developed to trace introgression patterns and to distinguish them from incomplete lineage sorting (Durand et al., 2011; Yu et al., 2012; Pease and Hahn, 2015; Martin et al., 2014). While these have been assessed for their power to identify introgression, they have the disadvantage that they require relatively rigid assumptions about the assumed history of introgression between them. These model assumptions are violated when introgression occurs from non-sampled sources, or repeatedly between the taxa. While such more complex scenarios could potentially be integrated into more formalized models, this would increase vastly the number of test comparisons in the statistics. In fact, there is no simple solution for this problem, implying that the procedure used to trace introgression need to be somewhat adapted to the taxa and the question that one wants to ask.

We have used here a framework of mouse populations and outgroups for which the phylogenetic histories are well known, based on fossil and phylogeographic evidence (Guénet and Bonhomme, 2003; Phifer-Rixey and Nachman, 2015; Hardouin et al., 2015). Figure 1 depicts these populations and their relationships.

**Figure 1.**
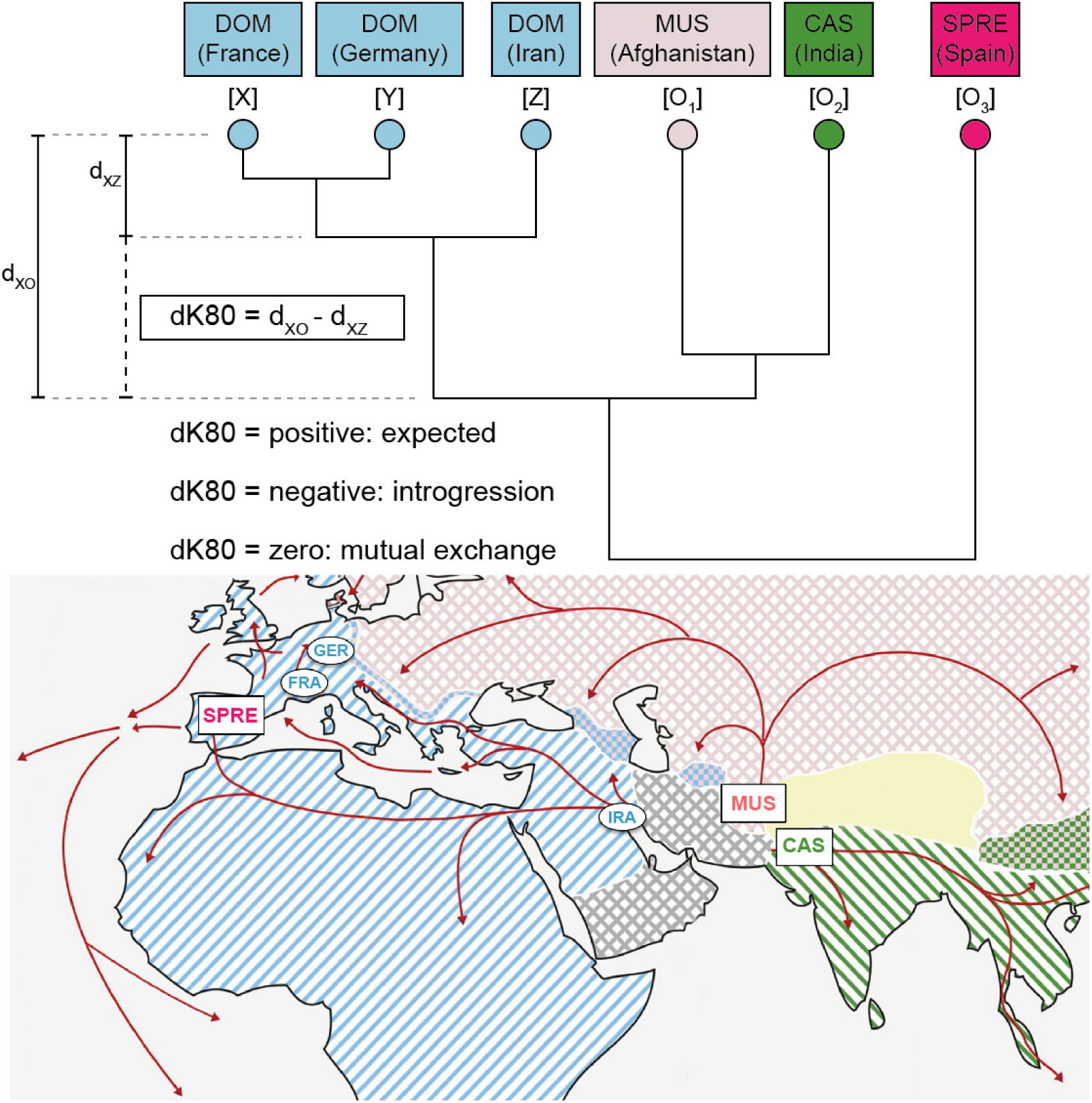
Relationships and origins of the mouse populations in the study. Three populations represent *M. m. domesticus* (DOM) (from France = Fra, Germany = Ger and Iran = Ira), the population from Afghanistan represents *M. m. musculus* (MUS), the one from India *M. m. castaneus* (CAS) and the one from Spain *M. spretus* (SPRE). The map shows the approximate locations, as well as the known dispersal routes (picture modified from Harr et al. (2016)). The tree represents the relationships. The principle of calculating dK80 is depicted to the left for one particular example, namely the comparison of the distance between Fra-Ira with the one between Fra-MUS, whereby Ira is the in-group [*Z*] and MUS the out-group [*O*_1_]. The two alternative outgroups used in this study are CAS [*O*_2_] and SPRE [*O*_3_].

Our focal *M. m. domesticus* populations come from France (Fra) and Germany (Ger). These are derived from a population from Iran (Ira), which invaded Western Europe about 3,000 years ago (Cucchi et al., 2005; Hardouin et al., 2015). The Fra and Ger populations have split shortly after arrival in Southern France and have since developed largely independently (Ihle et al., 2006; Teschke et al., 2008; Staubach et al., 2012). Hence, if a secondary introgression has occurred after the split, it should become visible in only one of the populations. While any population could serve as a source for secondary introgression, including the one from Ira (Hasenkamp et al., 2015), our focus in this study are the two subspecies *M. m. musculus* (MUS - represented by a population from Afghanistan) and *M. m. castaneus* (CAS - represented by a population from India), as well as the species *M. spretus* (SPRE - represented by a population from Spain) as possible source populations.

To study introgression, we use full genome sequence information and a branch-length comparison approach (Figure 1). This approach allows to detect the most prominent regions of unequivocal introgression. We first applied it to a previously identified introgression event at the amylase gene cluster and provide evidence that the introgression is likely related to a rescue of a pseudogenized *Amy2b* allele. However, we noticed also that the introgression region as a whole has a much more complex history. By assessing the whole genome for similar patterns, we found that regions exist that are subject to apparent mutual introgression and haplotype exchange between the hybridizing taxa. Most notably, many of these regions code for loci involved in adaptive and innate immune defense, in the defense against transposable elements and some appear to be involved in meiotic drive. Several affect parts of the olfactory and vomeronasal receptor clusters. Many, but not all are in regions of gene clusters with copy number variation. These findings suggest that mechanisms exist that allow the frequent exchange of genes involved in frequent adaptive processes between the taxa, even though most of them are regionally separated and/or hybrids are sub-fertile.

## Results

We have previously generated genomic re-sequencing data for multiple individuals for each of our focal populations (Harr et al., 2016). We have used these reads and created a consensus sequence representative for each population. This was done to ensure that only haplotypes with a frequency higher than 0.5 are represented. Hence, we are tracing introgression patterns that are either on the way to fixation or fixed. Our previous study (Staubach et al., 2012) had shown that the majority of recently introgressed haplotypes segregate only at low frequency, hence by focusing on the high frequency variants in the present study, we are tracing genomic introgression regions that have most likely been subject to recent positive selection in the respective populations (see modeling in Staubach et al. (2012)).

### Population-specific introgression patterns

To identify genomic regions of introgression, we used a sliding window approach (25kb per window) and generated a phylogenetic tree for each window. Linkage disequilibrium drops fast within 20kb in wild populations (Laurie et al., 2007; Staubach et al., 2012), i.e. by focusing on 25kb window sizes, we trace mostly relatively recent events that have not been subject to much recombination. We noticed that tree lengths can vary considerably along the chromosomes, which makes a simple dxy analysis for tracing tree incongruence as indicators of introgression less suitable. To compensate for this, we use the subtraction measure deltaK80 (dK80) depicted in Figure 1. The trees serve to calculate dK80 by subtracting the distance of the focal groups (either Fra or Ger) to the founder population (Ira) from the respective distances to the tested outgroups (MUS, CAS or SPRE). dK80 is expected to be positive when no introgression has occurred and negative when introgression from the tested outgroup has occurred. Based on simulations of our tree configuration without introgression, we find that dK80 values have a normal distribution (see q-q plots in Figure 1 - figure supplement 1).

The data for the dK80 values are provided in suppl. Table 1 (comparisons with MUS as outgroup [*O*_1_), suppl. Table 2 (comparisons with CAS as outgroup [*O*_2_]) and suppl. Table 3 (comparisons with SPRE as outgroup [*O*_3_]). We have also plotted these actual data distributions in q-q plots and find that they are less dispersed and more skewed than the ones of the simulations (suppl. Figure 1). Interestingly, the skews occur not only in the negative direction, but also in the positive one. In the following we focus the analysis on the negative side and come back to the skew on the positive side in the discussion.

dK80 values can be easily visualized as genome browser tracks allowing to recognize even complex patterns of introgression (see below). We have generated a custom track set on the UCSC genome browser named “wildmouse-introgression” that includes all data discussed here. Examples of these tracks are in the figures below.

We surveyed all negative outlier windows beyond a cutoff of 0.01% of the distribution of the real data (3.89 SD). Since about 100,000 windows were surveyed, one would expect around 5 by chance in a normal distribution, but we find between 99 to 352 in the different comparisons per population (Table 1; genome positions in suppl. Table 4).

**Table 1.** dK80 averages and outliers in the populations.

|  | tested outgroup population |  |  |
| --- | --- | --- | --- |
|  | MUS | CAS | SPRE |
| windows (N) | 102,281 | 102,299 | 102,009 |
| dK80 average (x103) | 6.08 | 4.15 | 14.94 |
| dK80 StDEV (x103) | 3.54 | 2.97 | 4.22 |
| 0.01% cutoff (x103) | -7.69 | -7.40 | -1.50 |
| smallest dK80 (x103) | -55.07 | -56.17 | -45.53 |
| windows (N) smaller than cutoff at 0.01% |  |  |  |
| total in Fra | 114 | 115 | 282 |
| total in Ger | 99 | 134 | 352 |
| total in Fra and Ger | 55 | 65 | 178 |
| only in Fra | 59 | 50 | 104 |
| only in Ger | 44 | 69 | 174 |

Among these we find between 55 - 178 in both, the Fra and the Ger population. This can be interpreted either that both populations were introgressed, possibly already in their direct colonizing ancestor, or that the Ira population has itself become introgressed from an unknown source after splitting from the ancestor of the Western European populations. Note that there are several further mouse lineages as potential donors in Iran that have not been characterized in detail yet (Hardouin et al., 2015). We have inspected the regions including outlier windows in the genome browser tracks and found that they represent often complex patterns of introgression. We first focus on a region coding for amylase genes.

### *Amylase 2b* introgression

We had previously identified the region around the pancreas Amylase 2b (*Amy2b*) as having been subject to adaptive introgression in the Fra population (Staubach et al., 2012). The dK80 approach and the outlier windows identify the same region, but the better resolution available through the full genome sequences allows now a much more detailed picture (Figure 2). The overall introgression (i.e. negative dK80 values) from *M. m. musculus* into Fra is as broad as originally found (approx. 0.5Mb), but the corresponding track lines for introgression from *M. m. castaneus* and *M. spretus* are more complex. A particularly strong introgression signal for all three outgroups, identified by outlier windows (Figure 2, bottom tracks), covers part of the non-coding region between *Amy2b* and its duplicated paralogs (Figure 2). Hence, the historical introgression dynamics at this locus appears to have been much more complex than originally anticipated. Note that there are also complex introgression patterns in the neighboring *Amy1a* gene (see data tracks “wildmouse-introgression”) that are not further discussed here.

**Figure 2.**
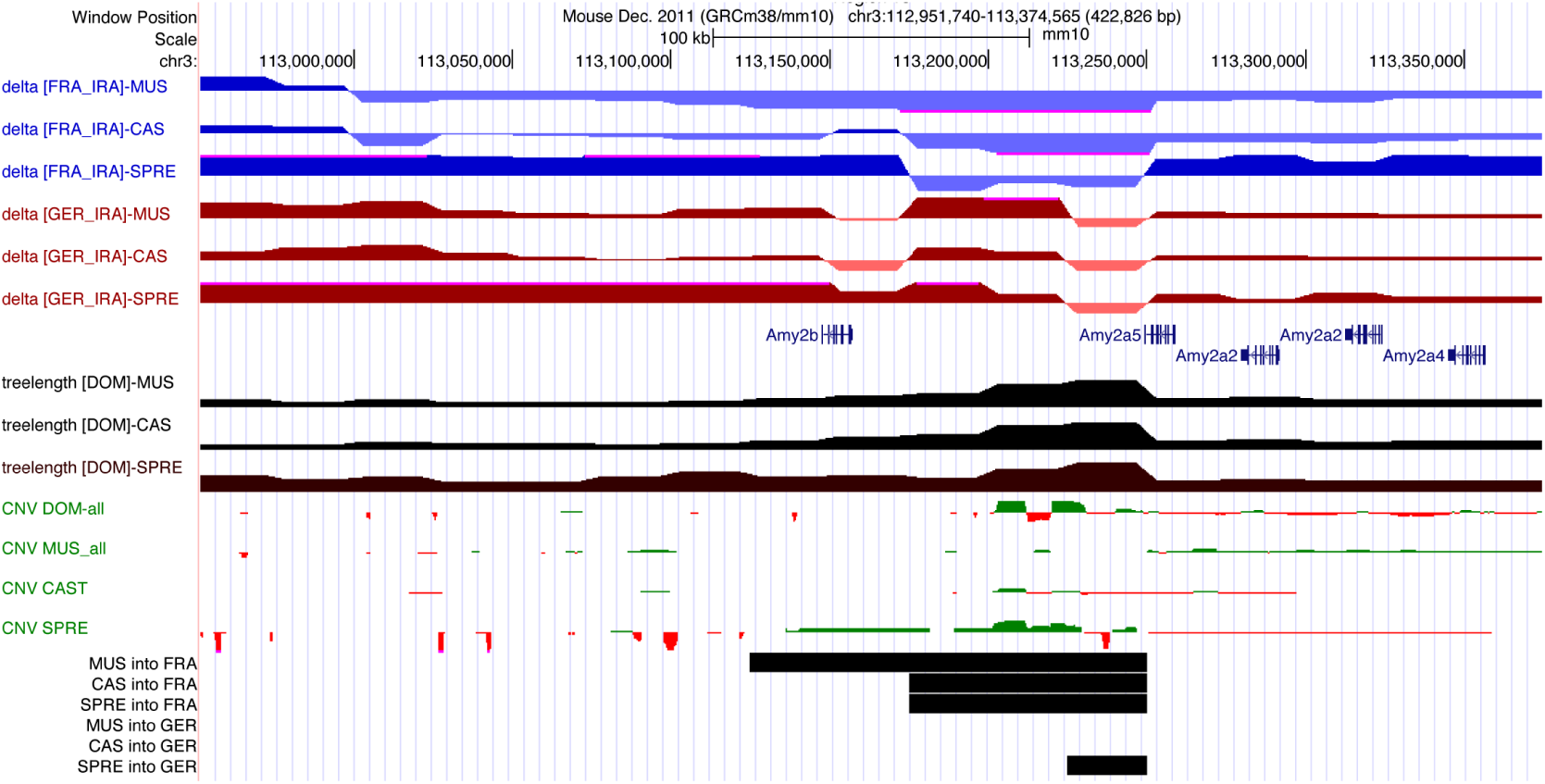
Introgression patterns around the amylase 2 gene cluster. Depiction of UCSC browser tracks for a region of chromosome 3 (positions in header). The rows show from top to bottom: the tracks for the dK80 values for the FRA comparisons (blue/magenta) and the GER comparisons (brown/red);gene annotations from the “UCSC Genes” track implemented in the UCSC Mouse Genome Browser (GRCm38/mm10); tree length tracks for the three outgroup comparisons (black);copy number variation tracks summed across all individuals of the respective populations, taken form Pezer et al. (2015) (green is more copies than reference, red is fewer copies than reference);windows with significant introgression in the indicated directions.

Interestingly, the sequence analysis provides a hint towards the possible reason for the apparently very recent introgression of a *M. m. musculus* haplotype into Fra. All of the Ger individuals sequenced harbor a mutation in the first exon that leads to a premature stop codon (Figure 3A). Hence, the Ger individuals carry a pseudogene for *Amy2b.* We have typed this variant for an extended sample of animals and populations and find the pseudogene variant to be prevalent in Germany, but rare in France (Figure 3b). The *Amy2b* sequences of the eight fully sequenced animals from Fra are all very similar and cluster closely with the *M. m. musculus* consensus sequence (Figure 3c). A native gel electrophoresis from pancreas tissue shows that there are still amylase variants in the Ger individuals, but the band patterns shows a composite between those found in the Fra animals and in *M. m. musculus / M. m. castaneus* animals. These are are most likely derived from the paralogs in the region, but a full resolution of the CNV haplotypes would be required to verify this. A quantitative activity assay shows that the overall amylase activity is lower in Ger than in Fra or *M. m. musculus* animals (Figure 3d). Hence, this pattern is compatible with the assumption that the mice colonizing Western Europe had lost the original *Amy2b* gene, but partially replaced this by activity from the duplicated copies in the vicinity. The Fra population has apparently captured a haplotype from *M. m. musculus* representing the originally active *Amy2b* gene. This haplotype has then spread rapidly in the Fra populations and is apparently now also invading Ger populations. Because an apparent rescue of an enzyme function is involved, this would have occurred through an adaptive spread, which explains also the selective sweep signature that we had detected in this region (Staubach et al., 2012).

**Figure 3.**
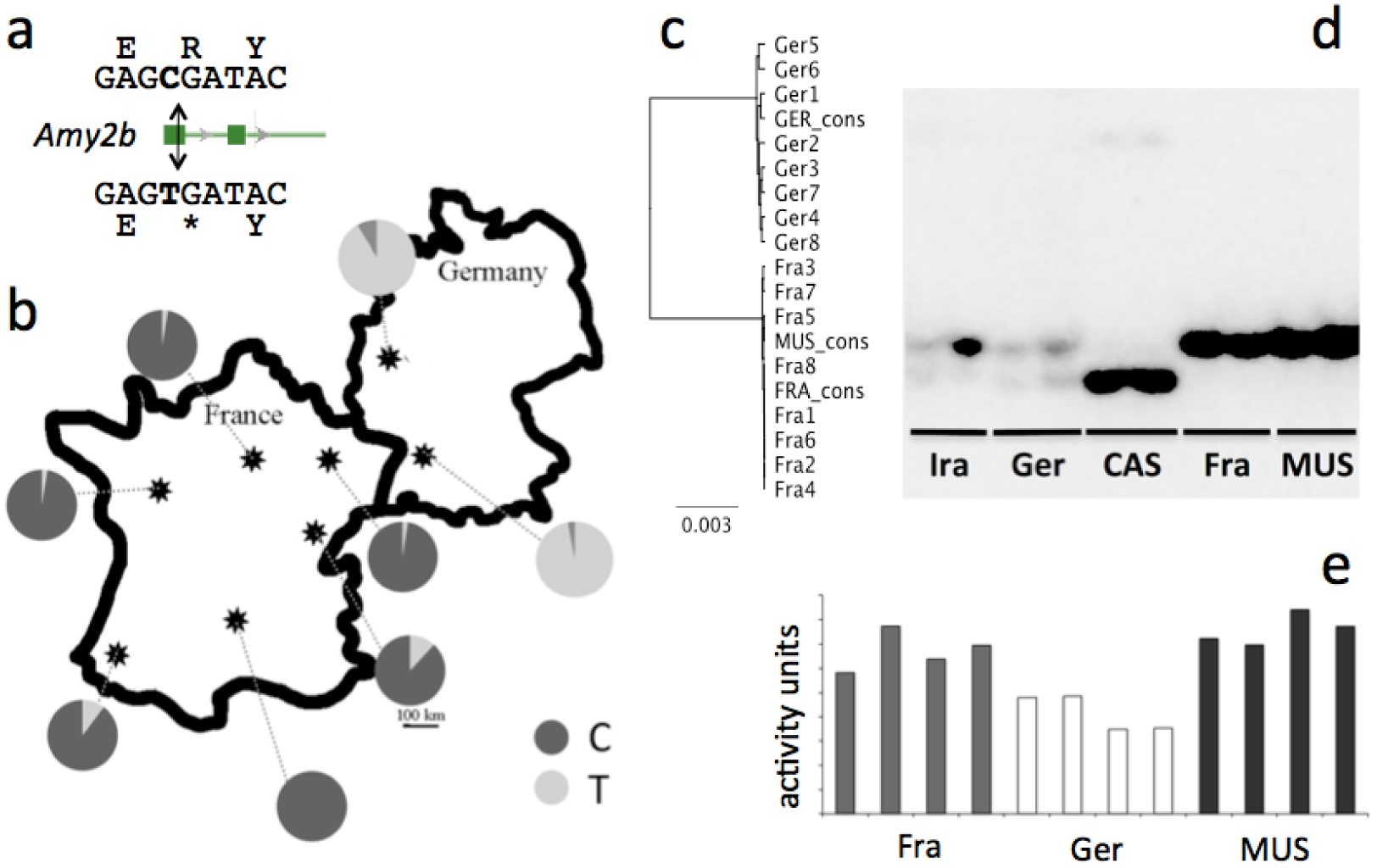
*Amy2b* introgression patterns. (a) Depiction of of the disabling mutation in exon1 of *Amy2b.* (b) Frequencies of the disabling (T-allele) versus enabling (C-allele) in different populations in France and Germany. (c) Phylogeny of *Amy2b* alleles of individuals and consensus sequences of the Fra and Ger population, plus consensus sequence of the MUS population. (d) Westernblot from a native gel of pancreas samples of two animals each of five populations, stained with an *Amy2b* antibody. Note that a denaturing gel does not resolve these variants (data not shown). (e) Amylase activity assays of four animals each from the three indicated populations.

The case of the *Amy2b* introgression presents thus a model of how one can envisage adaptive introgression taking place. An allele that has an advantage, either because it has itself acquired a new adaptation in a source population, or because it replaces a slightly deleterious mutation in the receiving population can apparently be specifically acquired, even from distant populations and can then reach high frequencies in the receiving population.

### Genomic introgression windows

The complexity of the introgression pattern shown in Figure 2 suggests that a region subject to introgression may actually be invaded repeatedly from different sources. If this would happen frequently, one would expect that dK80 would assume values around zero because the mutual invasions would equilibrate the distances between the respective groups.

We have therefore sought to systematically identify regions of such repeated introgression. However, because of the complexity of the possible patterns, they can not be easily captured by general statistical criteria. But the regions can readily be identified when one inspects the dK80 browser tracks. Since most of the genome follows the expected distribution of dK80 values, individual introgression regions become visible as noticeable dips in this background. Figure 4 shows two examples of 10Mb windows from chromosome 12 and chromosome 17. The chromosome 12 region includes an approximately 2Mb window with a complex mixture of strong introgression (identified by outlier windows) and flat distributions. The region codes for the mouse immunoglobulin heavy gene genes (*Igh*) encoding the exons that make up the variable antibody parts. In fact, another such region on chromosome 6 shows a similar pattern (see below). The chromosome 17 region includes an approximately 5Mb region that is relatively flat for comparisons between the sub-species *M. m. musculus* and *M. m. castaneus*, plus two shorter dips where also the comparison with M. spretus shows introgression, including significant outlier windows. These are enlarged at the bottom of Figure 4 and show that they code for key genes of the major histocompatibility complex (MHC), namely the antigen fragment binding receptors *H2*-*A*, *H2*-*B*, *H2*-*O* and *H2*-*Q.* Transspecies alleles have been described for *H2*-*A* and *H2*-*B* genes before, supporting the notion that our procedure picks out such introgression regions faithfully.

**Figure 4.**
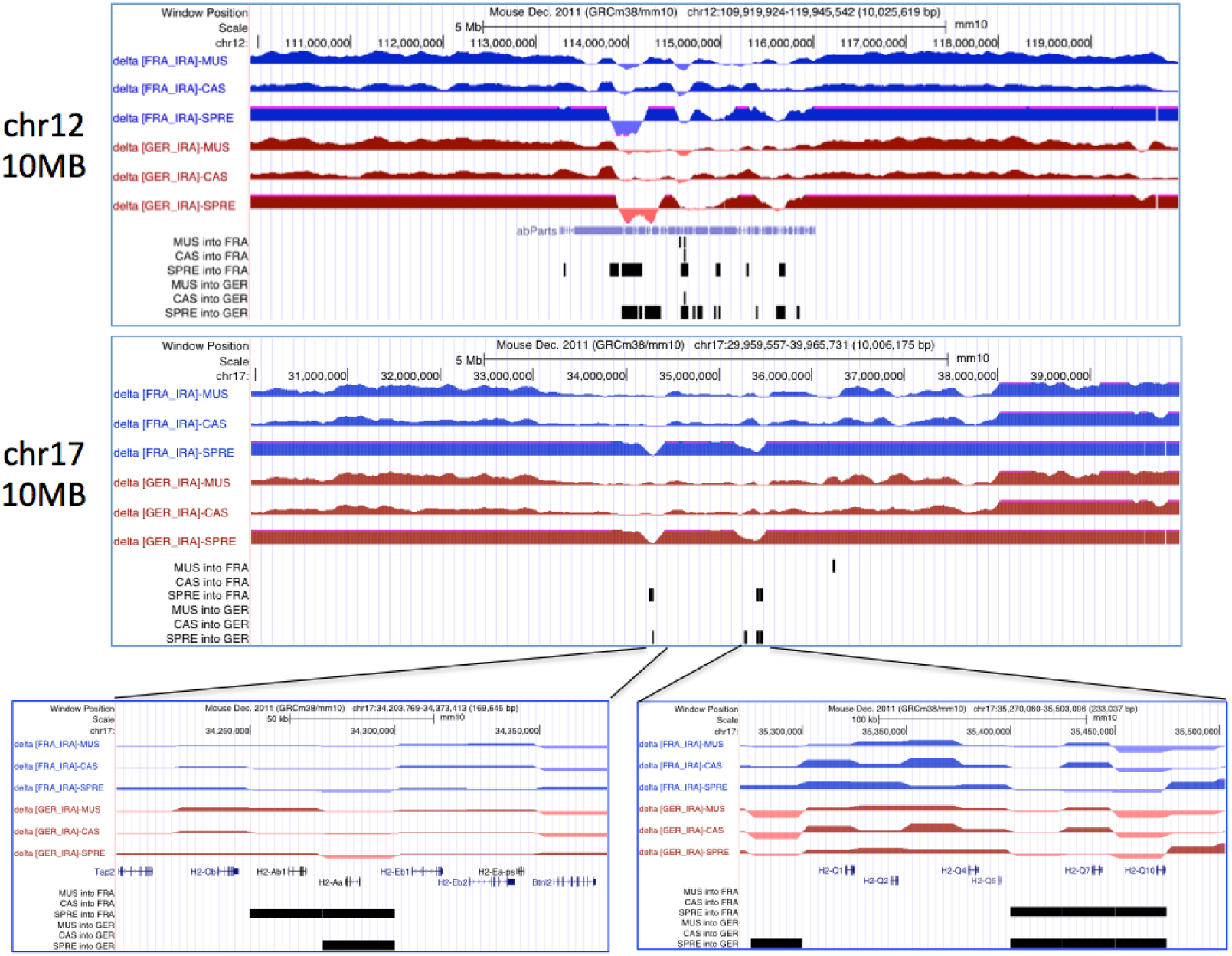
Two examples of mutual introgression windows. Depiction of UCSC browser tracks for 10MB regions of chromosome 12 and chromosome 17 (positions in headers). The rows show from top to bottom: the tracks for the dK80 values for the FRA comparisons (blue/magenta) and the GER comparisons (brown/red); gene annotations from the “UCSC Genes” track implemented in the UCSC Mouse Genome Browser (GRCm38/mm10); windows with significant introgression in the indicated directions.

We have inspected all 10MB windows across the whole genome (excluding the Y-chromosome because of too many missing data) to identify a set of windows that show unusual patterns across all three outgroup comparisons. Table 2 lists all of these regions with a length of at least 100kb (i.e. 4 consecutive windows). The majority (12 out of 18) have well defined functions in adaptive immunity (5), innate immunity (7) and three represent KRAB Zn-finger genes which function in the repression of transposable elements Jacobs et al., 2014; Imbeault et al., 2017; Kauzlaric et al., 2017). Two cover parts of olfactory receptor gene clusters whereby region 9 includes besides an olfactory receptor cluster also the hemoglobin beta-chain genes, which are known to be subject to complex selection and gene replacement in mouse (Storz et al., 2007). The track patterns for the 18 regions listed in Table 2 are compiled in Figure 4-Figure supplement 1.

**Table 2.**
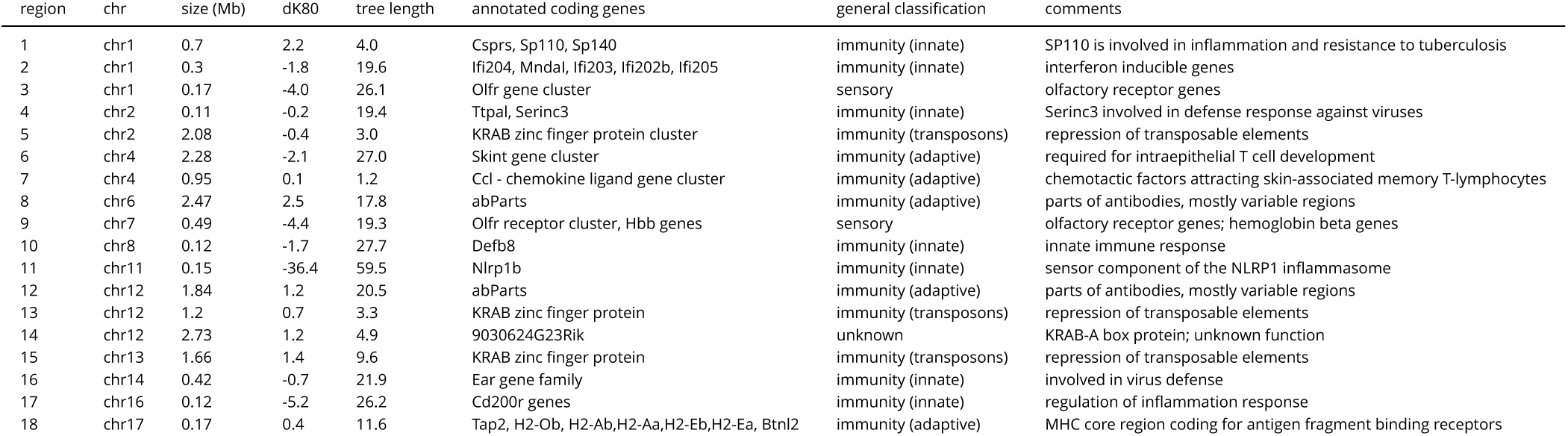
Introgression between all in-groups and out-groups. #chromosomal genome positions are provided in Figure 4-Figure supplement 3; dK80 and tree length represent averages ×1000; “immunity” is classified into three categories: “adaptive” = adaptive immunity, “innate” = innate immunity and “transposons” = transposon repression

In addition to the regions involving mutual introgression with *M. spretus*, we have also systematically surveyed all regions >100kb that involve mostly exchange between the three sub-species (*M. m. domesticus*, *M. m. musculus* and *M. m. castaneus*) by the same criteria. These include a total of 67 regions (suppl. Table 5), including the overlaps with the ones listed in Table 2. 29 of the regions have clear immune functions, eight represent olfactory and vomeronasal receptor clusters and another eight represent testis specific genes, or genes highly expressed in testis. Six regions each cover non-coding parts of the genome (i.e. no annotations in the respective window) and genes with unknown functions (suppl. Table 5).

Inspection of the windows shows that tree lengths differ much in the different regions. Some introgression regions have very long trees, others very short ones (Table 2 and suppl. Table 5). On average, there is a good correlation between tree length and dK80 score (r2 = 0.69 for the data in Table 2 and 0.45 for the data in suppl. Table 5), with more negative scores showing the longer trees. The most negative score combined with the longest average tree is found for *Nlrp1b*, which codes for the sensor component of the inflammasome (region 11 in Table 2). This pattern is apparently caused by an introgression of a highly unrelated haplotype into the Ira population (tree in Figure 4-supplement 2), which renders all other comparisons negative. The shortest average tree length is found for the window coding for a chemokine ligand cluster (region 7 in Table 2), rendering this 0.92Mb region highly similar between all analyzed taxa (tree in Figure 4-supplement 2). This suggests that a given haplotype variant has introgressed into all of them, possibly located on an inversion, since the size appears to be the same in all taxa.

The gene clusters identified within the windows show often copy number variation (see suppl. Figure 4-Figure supplement 1). Of the three single genes with major copy number variation in natural populations identified in Pezer et al. (2015), two show patterns of introgression. One is *Cwc22*, which encodes a splicing factor. Its introgression pattern is apparent in the comparison with CAS and SPRE (Figure 5a). The other is *Hjurp*, which codes for a holiday junction recognition protein. For this gene, introgression is mostly evident for the Ger population with respect to MUS and CAS introgression (Figure 5b).

**Figure 5.**
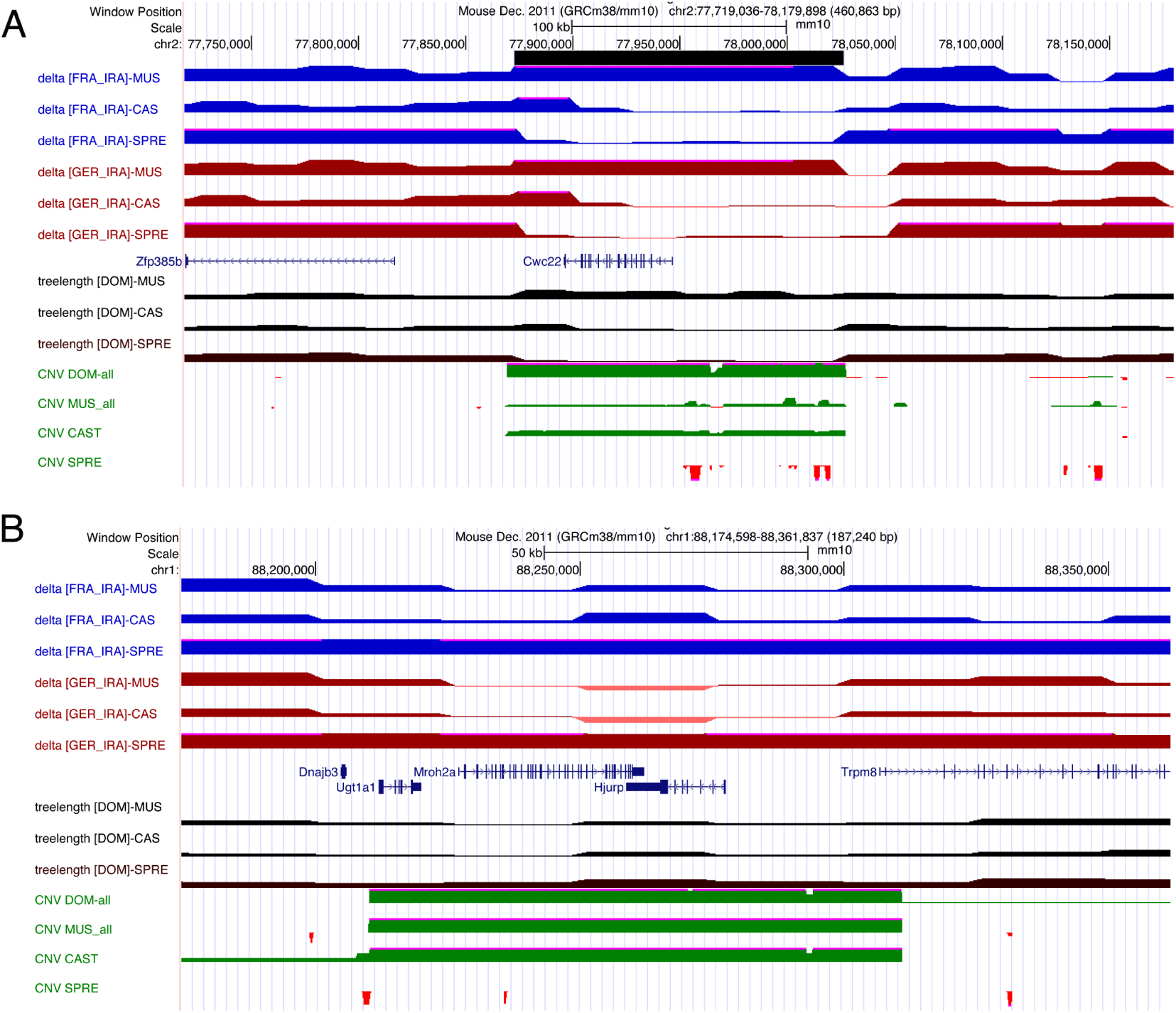
Introgression patterns in highly copy number variable regions. Depiction of UCSC browser tracks for chromosomal regions of *Cwc22* (A) and *Hjurp* (B) (positions in headers). Rows from top to bottom as in Figure 2. *Cwc22* and Hjurp were previously found to be among the most copy number variable genes between mouse populations (Pezer et al. 2015). *Cwc22* shows introgression in comparison to *M. m. castaneus* and *M. spretus*, for *Hjurp*, the pattern is most evident for the comparison to *M. m. musculus* and *M. m. castaneus* in the Ger population.

## Discussion

Our initial data analysis was geared towards identifying specific regions of introgression in populations that have established only a few thousand years ago (Fra and Ger). One such region that we had identified before is a region on chromosome 3 coding for an amylase gene cluster. Here we confirmed this region, based on a procedure specifically developed to identify regions of potentially adaptive introgression. In the case of the amylase region, we show that a secondary replacement of pseudogenized version of *Amy2b* was apparently the reason for the introgression in one of the *M. m. domesticus* sub-populations. However, our analysis revealed that the patterns are much more complex. There are apparently many regions in the genome that are subject to repeated introgression in different directions, with apparently different histories, and including haplotypes from sources that have not been sampled. Given the abundant evidence for introgression in many studies, this should not be so surprising. In fact, this complexity makes average genome-wide statistical analyses complicated, since these can only work within the framework of given scenarios and do not account for repeated or mutual introgression. Our study has initially also focused onto a particular scenario, namely introgression into newly established populations from other sub-species and species. But the fact that many of the outlier windows identified through this procedure overlap and cluster in specific regions revealed an underlying complexity that could eventually only be resolved through direct inspection of introgression patterns.

By using consensus sequences from the populations, our approach is conservative, since we miss out cases where an introgressing haplotype is still at a frequency below 50%. This is for example the case for the otherwise well supported *Vkorc1* region that was suggested to be in a phase of adaptive introgression in some Western European populations (Song et al., 2011; Liu et al., 2014). In our data, we find only a subset of animals in the Fra and Ger populations carrying this haplotype, i.e. it does not show up among the consensus sequences.

Our focus was on low or negative dK80 values, since these would indicate introgression within our general tree topology. However, the q-q plots in Figure 1-Figure supplement 1 show that deviations from the expected distributions occur also at high dK80 values. We have inspected these outliers and find that they represent another level of complexity, such as introgression into the outgroup population from unknown sources, making the distances much larger than average. It is known that there are further sub-species or species in Asia (Hardouin et al., 2015; Hamid et al., 2017) that could be the source for such introgression. Given our general finding of mutual introgression among our sampled populations, it is not unexpected that this should also occur with non-sampled populations.

### Introgression in evolutionary active regions

The inspection of the dK80 tracks across the whole genome and across the different populations allowed to pinpoint regions of particularly active introgression. Intriguingly, the majority of these regions is related to adaptive and innate immune functions. This includes the MHC, where transspecies alleles are often found (Parham et al., 1996). These have usually been ascribed to be the result of balancing selection and/or incomplete lineage sorting after the splitting of the species. However, for species that have remained at least partially inter-fertile, there is increasing evidence that introgression must play a role as well (Abi-Rached et al., 2011; Grossen et al., 2014). This is now fully confirmed in our study. Key genes of the core MHC region, including H2-Aa, H2-Ab, H2-Ea and H2-Eb show significant introgression windows for the comparisons with *M. spretus.* Further, an extended 2.9Mb region shows a flat dK80 pattern for the comparisons with the subspecies (Figure 2). These regions are too long to be considered as remnants from the progenitor populations maintained by balancing selection, since they would have been broken up by recombination. Interestingly, in spite of the apparent frequent introgression, at least the loci coding for the H2-Aa, H2-Ab receptors show long trees, which is compatible with balancing selection in this region to maintain a high diversity, which may even be fueled by capturing alleles from other sub-species and species.

While the introgression pattern within the MHC is not unexpected, it had remained so far unnoticed that many other immune genes are also subject to introgression. Most notably, these encode the antibody coding regions on chromosomes 6 and 12, which are also part of the adaptive immune system. Since the antibody diversity is generated through splicing among variable exons, it seems that this diversity is also enhanced by taking up alleles from other populations.

Another immune region with strong introgression and long trees is the *Skint* gene family cluster on chromosome 4 (region 6 in Table 2). *Skints* code for proteins containing trans membrane spanning domains and extracellular IgV and IgC domains that are specifically expressed in dendritic epidermal T-cells which play a crucial role in immune defense during wound healing (Keyes et al., 2016). Interestingly, a second introgression region on chromosome 4 (region 7 in Table 2) codes for a cluster of cytokine genes including *CCL27*, which has been shown to selectively recruit cutaneous memory T lymphocytes into the skin (Morales et al., 1999) and is generally implicated in regulating wound repair (Hocking, 2015). This region shows, however, a very flat signature, both for dK80, as well as for tree length, indicating mutual introgression or a recent sweep of a whole haplotype across the tested species and sub-species (Figure 4-Figure supplement 2. A third introgression region associated with epidermis is the one that encodes the *Defb8* gene (region 10 in Table 2). ß-defensins code for antimicrobial and chemo-attractant peptides, especially attracting CD4+ T-cells (Taylor et al., 2009) and high expression of *Defb8* occurs in epidermal tissues.

Three introgression regions (region 1,11 and 17 in Table 2) are involved in inflammation regulation. This includes the *SP110* gene that is involved in resistance to tuberculosis (Wu et al., 2016), *Nlrp1b*, a sensor component of the inflammasome (Chavarría-Smith and Vance, 2015) and *CD200r* genes regulate macrophage function in inflammation reactions (Snelgrove et al., 2008; Fraser et al., 2016).

Another three regions code for genes that are interferon inducible and/or involved in virus defense. These include a cluster of interferon inducible genes on chromosome 1 that function in virus recognition (*Ifi203* - region 2) (Stavrou et al., 2015), activity against retroviruses (*Serinc3* - region 4) (Usami et al., 2015) and influenza virus protection in epithelia (*Ear1* - region 16) (O’Reilly et al., 2012). The fact that all of these regions are similarly, or even more strongly affected by introgression than the MHC genes suggests that they play a similar major role in counteracting fast evolving pathogens. Hence, the current focus on MHC genes as the major driver of evolutionary responses to pathogens may be too limited. It seems warranted to pay additional attention to immune processes in the epithelial cells, as well as innate mechanisms of virus defense. Differences in innate immune responses in human populations have also been ascribed to adaptive introgression of Neandertal alleles (Quach et al., 2016).

Immunity is not only relevant against pathogens, but also against transposable elements. KRAB zinc finger proteins have been implicated in this function, whereby there is an evolutionary arms race between adaptation of the zinc fingers to the recognitions sites in the transposable elements and the acquisition of new mutations in the recognition sites (Jacobs et al., 2014). We find a total of four clusters of such genes plus a single gene one among the introgression regions (suppl. Table 4). One of them (region 13 - Table 2) covers a 2Mb region with almost identical sequences between all taxa (i.e. very short trees). In fact, one should expect that active transposable elements could introgress as well between the taxa, i.e. it makes sense when they share the same set of defense genes.

The extended table with introgression regions that show a signal with only two of the three outgroups (suppl. Table 5) provides some further interesting insights into the type of processes affected most by introgression in mice. There are six regions that code for olfactory receptor clusters, some as specific parts of larger such clusters and two of the several vomeronasal receptor clusters in the genome show an introgression signal. This would suggest that they code for sub-types that may be particularly relevant for an evolutionary active process.

There are four regions that code for SPEER/Takusan domain genes. These were originally identified as testis-specific genes (Spiess et al., 2003) and they occur in several clusters in the genome. The genes have specifically evolved in the rodent lineage, possibly through some protein domain fusions and fast subsequent evolution. The variants occurring on chromosome 14 were also implicated in the regulation of synaptic activity (Tu et al., 2007) but these are also more expressed in testis than in brain (see also expression data provided in Harr et al. (2016)). In their study on postmeiotic transcription in sperm cells, Moretti et al. (2016) speculate that these genes may act in a meiotic drive dynamics, possibly as suppressors of the intragenomic conflict between *Slx* and *Sly* gene families (Helleu et al., 2015). Interestingly, the *Slx* gene cluster on the X chromosome shows also corresponding introgression patterns (suppl. Table 5) and, the *Sly* gene cluster on the Y-chromosome shows similar patterns (although we have not included these in our analysis because of too many missing data, we provide the partial data in the track patterns of “wildmouse-introgression”).

Another region that was suggested to be involved in transmission ratio distortion is *Cwc22* or *R2D2* on chromosome 2 (Didion et al., 2016). Population genetic studies of this region have shown that it shows major copy number variation changes between populations (Pezer et al., 2015; Didion et al., 2016) and that it was involved in recent selective sweeps (Didion et al., 2016). Hence, this is also a region with clear introgression patterns that has a proven high evolutionary dynamics.

## Conclusions

Although the populations we have included in our analysis are all very distinct, either due to allopatry or long evolutionary separation, they exchange nonetheless genes. Most of the genes that are exchanged have evident adaptive significance, including the case of the *Amy2b* gene, where a pseudogene allele became replaced by an active allele from another subspecies. Given that our study has focused on high frequency variants and that most of them are associated to various forms of immune defense, we can assume that most of the introgression events that we are tracing here have been adaptive. Hence, by identifying these introgression regions, we find at the same time candidate genes for frequent adaptations in mice.

## Methods and Materials

### Ethic statement

Mice were caught as described in Harr et al. (2016). Transportation of live mice to the animal facility, maintenance and handling were conducted in accordance with German animal welfare law (Tierschutzgesetz) and FELASA guidelines. Permits for keeping mice were obtained from the local veterinary office ‘Veterinäramt Kreis Plön’ (permit number: 1401-144/PLÖ-004697).

### Data and mouse samples

Genome data used in this study (*M. m. domesticus* GER - 8 individuals, *M. m. domesticus* FRA - 8 individuals, *M. m. domesticus* IRA - 8 individuals, *M. m. musculus* AFG - 6 individuals, *M. m. castaneus* CAS - 10 individuals, and *M. spretus* SPRE - 8 indiciduals) were taken from Harr et al. (2016). Mouse samples were taken from the collection at our institute - sources are described in Harr et al. (2016) and Linnenbrink et al. (2013).

### dK80 calculation

Quartets were used for all calculations of dK80. They included always the three DOM populations (Ger [*X*], Fra [*Y*] and Ira [*Z*]), as well as one of the three outgroups each, MUS [*O*_1_, CAS[*O*_2_] or SPRE [*O*_3_] (compare Figure 1). Within the quartets, dK80 was calculated for trios (either with Ger or Fra) on non-overlapping sequence windows (*w*) throughout the genome between this population triplet on a window (*w_i_*) as:

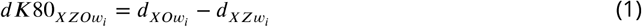

where *d_XOw_i__* and *d_XZw_i__* are defined as the average Kimura’s 2-parameter sequence distance (Kimura, 1980) between the corresponding two populations calculated with the function ‘dist.dna’ of the R package ‘ape’ (Paradis et al., 2004) using the model ‘K80’. Prior the calculation of dK80 all sites with missing data (see below) within the specified window (*w_i_*) and the specified populations ([*X*], [*Z*], [*O*]) were removed across the whole quartet with the ‘Biostrings’ R package (Pages et al., 2009) and only those windows retained with a missing rate lower than 50%.

### Population specific masking

Sequences with low coverage (missing data) were excluded from the analysis by masking them for each quartet under analysis. Masking files were generated by processing the BAM files with ‘enomeCoverageBed’ and ‘mergeBed’ from the bedtools2 software suite (Quinlan and Hall, 2010, v2.26.0) to obtain site specific genome coverage per individual. Sites with a coverage smaller than 5 were used as masking regions for each individual. Further, to obtain population specific masking regions, site specific genome coverage files were united with ‘unionBedGraphs’ and sites with a united coverage smaller than 5 were used as masking regions for the populations. Suppl. Tables 1-3 provide for each window the numbers of masked (“missing”) sites. Windows that had more than 50% sites missing were not included in the overall analysis.

### Consensus sequences

To construct individual and population specific consensus sequences, we conducted first a SNP calling on the mapped BAM files taken from Harr et al. (2016). Briefly, BAM files for individuals grouped by population were processed with ‘samtools mpileup’ and ‘bcftools call’ (Li, 2011, v1.3.1) with relaxed mapping quality options (samtools mpileup: -q 0 -Q 10 -A -d 99999 -t DP,AD,ADF,ADR -uf mm10.fasta; bcftools call: -f GQ -m -v) to also retain information in CNV regions. We corrected also for individual sex and ploidy (chrY F 0; chrX M 1; chrY M 1; chrM F 1; chrM M 1). We note that many of the here described introgression regions occur in copy number variable regions, which are often filtered out in standard analyses that constrain the data to high quality mapped reads. In our simulations we paid particular attention to such regions to assess whether the inclusion of low quality mapped reads would lead to artifacts with respect to introgression signals, but we did not find this to be the case. If anything, the inclusion of these reads would only make distance larger rather than smaller.

The resulting population specific VCF files were recoded with ‘vcftools’ (Danecek et al., 2011, v0.1.15) to remove indels and to only retain variant sites (vcftools: -recode -remove-indels -recodeINFO-all -non-ref-ac-any 1) either for the complete population (-keep pop) or for the individuals (-keep ind). Further, the recoded population specific VCF file containing multiple individuals was parsed with ‘vcfparser.py mvcf2consensus’ (https://gitlab.gwdg.de/evolgen/introgression; -cdp 11) to obtain a CONSENSUS VCF file for each population. In brief, the allelic depth information for the reference and alternative allele was summed over all individuals per population by simultaneously removing all sites which had a total depth (reference plus alternative allelic depth) of smaller than 11 in the population (DP<11).

The CONSENSUS VCF was used in combination with the population specific masking region to construct chromosome alignments with the reference (GRCm38/mm10) and the python script ‘vcfparser.py vcf2fasta’ (https://gitlab.gwdg.de/evolgen/introgression; -type refmajorsample -R mm10.fasta -cov2N 4) to obtain the major allele per variant and to mask all regions with a coverage smaller than 5. This step was repeated for individuals using the individual masking regions resulting in CONSENSUS and INDIVIDUAL FASTA files used for the inference of introgression.

### Simulations

Simulations were performed to evaluate dK80 under a phylogenetic scenario mimicking the real data without introgression, as well as to ensure that the mapping procedure does not lead to artefacts, especially in the copy number variable regions (see Figure 1-Figure supplement 6 for the simulation design).

First, to provide the appropriate distance framework, all pair-wise polymorphic sites and all pair-wise informative sites (excluding pair-wise missing sites) of all autosomes were counted between the investigated populations using the CONSENSUS FASTA files. The resulting pair-wise distances defined as:

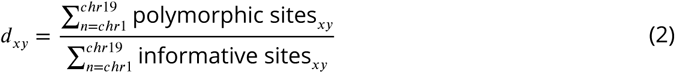

where *d_xy_* is the pair-wise distance between the consensus sequence of population *x* and *y*, were then used as a distance matrix to calculate an “Unweighted Pair Group Method with Arithmetic Mean” (UPGMA) tree in R. The resulting UPGMA tree distances were used as a proxy to simulate chromosome 1 with the python script ‘simdiv.py’ taking the phylogenetic context into account.

Second, the simulated sequences representative of each population (SPRE, CAS, MUS, Ira, Ger, Fra) were used to generate artificial Illumina reads to test the influence of possible sequencing errors with ‘ART’ (Huang et al., 2011, v2.5.8) and to mimic the original sequence libraries (art_illumina: -sam -na -ss HS20 -f 20 -l 100 -m 250 -s 80 -p). Subsequently, the artificial Illumina reads were mapped against the simulated reference with ‘bwa mem’ (Li and Durbin, 2009, 0.7.12-r1039), followed by sorting, marking and removing duplicates with the picard software suite (https://broadinstitute.github.io/picard/) and an indel realignment step with ‘GATK’ (McKenna et al., 2010, v3.7) as described in Harr et al. (2016). Masking, SNP calling, FASTA sequence construction and dK80 calculation was performed as described above.

### Data visualization and availability

Data visualization was done with the UCSC genome browser for the mouse assembly mm10. Custom tracks were generated and and are available as “Public Session” under the term “wildmouse-introgression”. Individual and population specific consensus sequences can be accessed via ftp http://wwwuser.gwdg.de/evolbio/evolgen/wildmouse/introgression/.

### Amylase purification and quantification

Approximately equal weights of pancreas per sample were used for purification. Tissues were homogenised in PBS using a TissueLyser II (Qiagen), centrifuged at 13,000 × g at 4°C for 10 minutes, and the crude lysate was collected. Ethanol was added to a final concentration of 40%, centrifuged at 10,000 × g for 10 minutes at 4°C, and the supernatant was collected. Amylase was precipitated by addition of 1mg of oyster glycogen (Sigma-Aldrich) according to Schramm and Loyter (1966), followed by shaking on ice for 5 minutes. It was then pelleted by centrifugation at 5,000 × g for 3 minutes at 4°C. The samples were washed, re-suspended in PBS, and glycogen digested by incubation at 30°C for 20 minutes (Hjorth, 1979). Samples were stored at −80°C in aliquots to avoid repeated freeze/thaw cycles.

Protein concentration was determined using Thermo Scientific’s Coomassie Plus^™^ (Bradford) Assay kit according to the manufacturer’s instructions as follows. A standard curve was generated using the BSA provided at 2000,1500,1000, 750, 500, 250,125 and 25*μ*g/mL. All unknown samples were diluted 1:2, and 10*μ*L of each sample and standard was added to a 96 well plate in duplicate. A PBS blank was also included. 300*μ*L of Coomassie Plus Reagent was added to each well and mixed on a plate mixer for 30 seconds. The plate was incubated at room temperature for 10 minutes, and then measured on a Tecan Infinite^®^ M200 PRO plate reader at 595nm. The blank measurement was subtracted from all other readings, and then the concentration of the unknown samples determined using the standard curve, and multiplied by the dilution factor.

### Native PAGE

Amylase extracts were separated on 7.5% Mini-PROTEAN^®^ TGX^™^ gels (Bio-Rad), but in their native form (no boiling, no SDS). Gels were loaded with 1.5*μ*g of each sample (using Bradford assay measurements) and run for 45 minutes at 100V, followed by 3 hours 45 minutes at 300V, with the tank immersed in ice throughout (adapted from Hjorth, 1979). Western blot analysis was done by semi-dry transfer to PVDF membrane for one hour at 20V, followed by blocking the membrane with 5% milk powder in PBS supplemented with 0.1% tween for one hour at room temperature. The membrane was probed for amylase using anti-*α* -amylase antibody from Cell Signalling (#4017) at 1:2000 in 2% milk in PBS tween overnight at 4°C. The primary antibody was detected using goat anti-rabbit HRP (Southern Biotech) at 1:5000 for one hour at room temperature followed by incubation with Immun-Star^™^ WesternC^™^ Chemiluminescent substrate (Bio-Rad). Blots were visualised using an Alpha Innotech FluorChem^™^ MultiImage^™^ light cabinet.

### Amylase activity

The activity of each sample was determined using an amylase activity assay kit (Sigma-Aldrich), which measures a colorimetric product resulting from the cleavage of ethylidene-pNP-G7 by amylase to generate p-nitrophenol. Samples were measured with a slightly modified version of the manufacturer’s instructions as follows. A nitrophenol standard curve was generated by adding 0, 2, 4, 6, 8, and 10*μ*L of 2mM Nitrophenol Standard to a 96 well plate and adding water to a final volume of 50*μ*L. Unknown samples were diluted 1:5000 and 50*μ*L added to the plate in duplicate. Amylase Assay Buffer and Amylase Substrate Mix were mixed together 1:1 and 100*μ* L added to each well. After 3 minutes incubation (= *T_initial_*) a reading was taken at 405nm (*A*405_*initial*_). The plate was incubated at 25°C and the absorbance measured every 2 minutes (as opposed to every 5 minutes in the manufacturer’s instructions). Readings were taken until the value of the most active sample exceeded the linear range of the standard curve. *A*405_*final*_ was taken as the penultimate reading before this. The background was subtracted, and the change in absorbance from *T_initial_* to *T_final_* calculated: Δ*A*405 = Δ*A*405_*final*_ - *A*405_*initial*_. The amount of nitrophenol generated between *T_initial_* to *T_final_* was calculated using the standard curve. The activity was then determined using the equation:

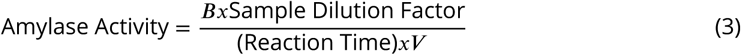

where: *B* = Amount (nmole) of nitrophenol generated between *T_initial_* and *T_final_* Reaction Time = *T_final_* - *T_initial_* (minutes) *V* = sample volume (mL) added to well One unit of amylase is the amount of amylase that cleaves ethylidene-pNP-G7 to generate 1.0 m mole of p-nitrophenol per minute at 25°C.

## Acknowledgments

We thank Emre Karakoc for the initial exploration of algorithms to detect introgression patterns in the mouse dataset, Ellen McConnell for the alpha-amylase experiments and the members of our lab for comments on the manuscript.

## Additional information

### Competing interests

DT: Senior editor, *eLife.* The other authors declare that no competing interests exist.

---

**Figure 1–Figure supplement 1.**
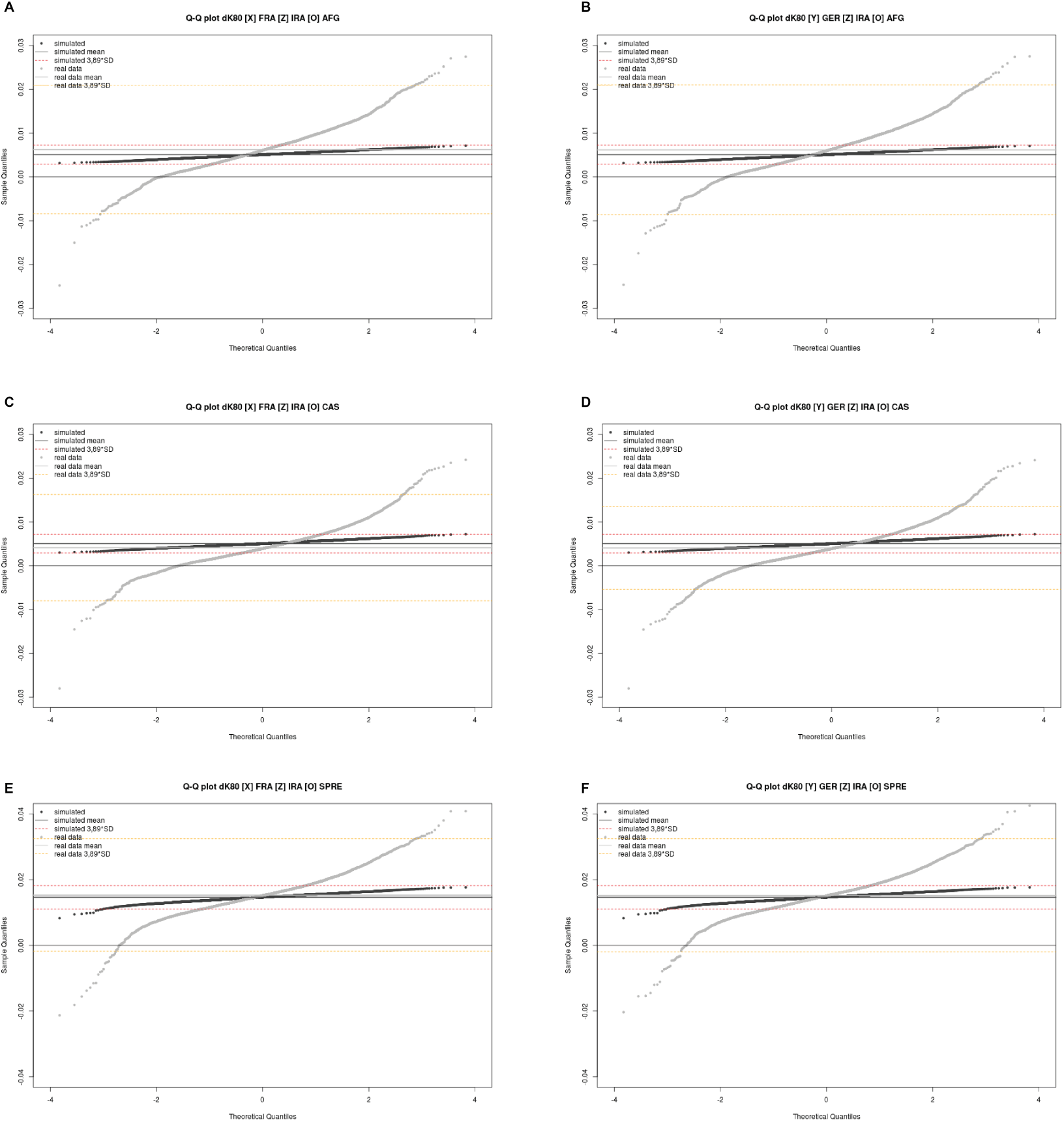
dK80 q-q plots for different population combinations. (A) [*X*]: Fra [*Z*]: IRA [*O*_1_]: AFG, (B) [*Y*]: Ger [*Z*]: IRA [*O*_1_]: AFG, (C) [*X*]: Fra [*Z*]: IRA [*O*_2_]: CAS, (D) [*Y*]: Ger [*Z*]: IRA [*O*_2_]: CAS, (E) [*X*]: Fra [*Z*]: IRA [*O*_3_]: SPRE, (F) [*Y*]: Ger [*Z*]: IRA [*O*_3_]: SPRE. Mean of dK80 distribution is highlighted as solid lines (black: simulated data; grey: real data), 3.89 standard deviations are highlighted by dashed lines (red: simulated data; orange: real data).

**Figure 1–Figure supplement 2.**
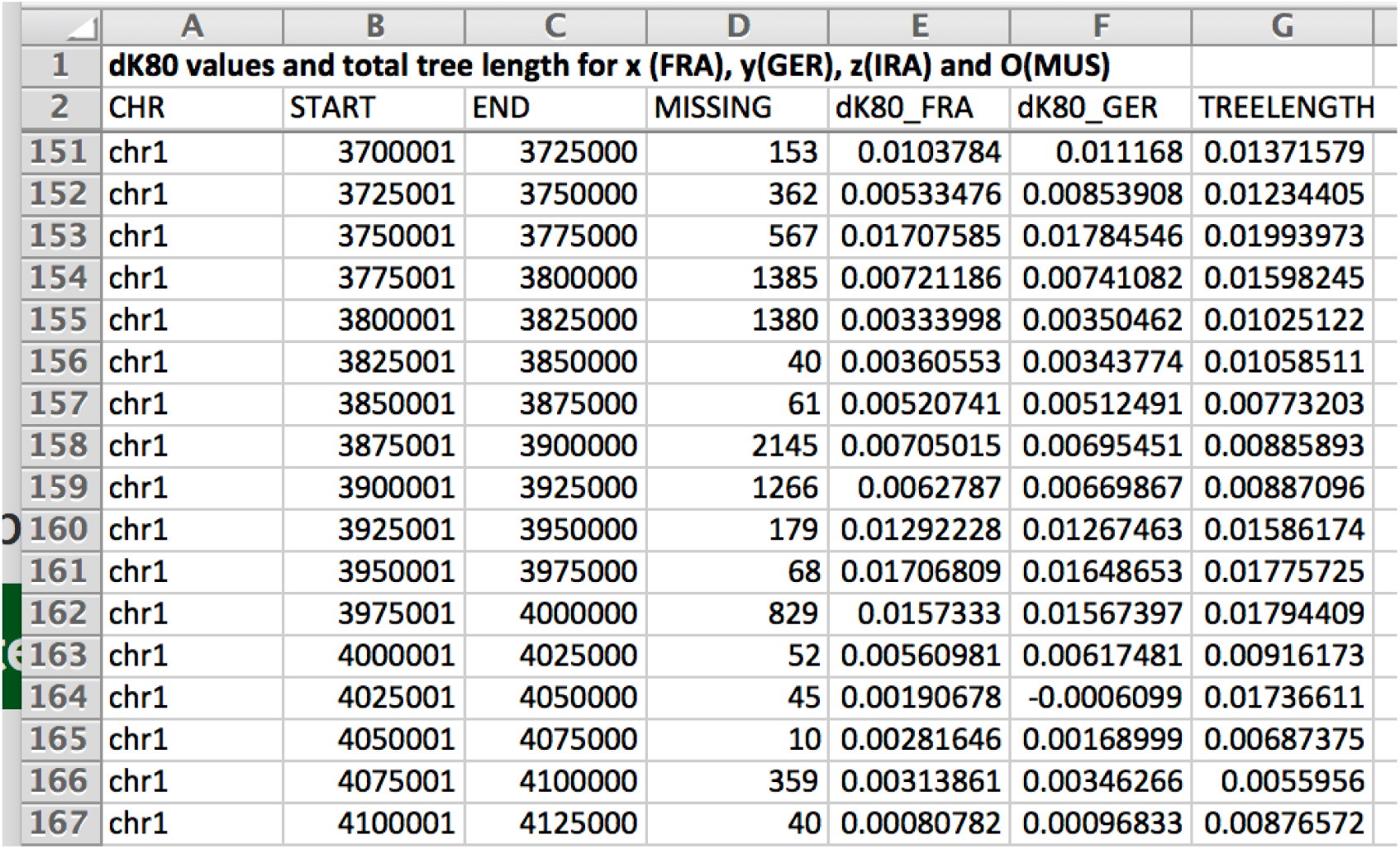
Preview of supplementary Table 1. dK80 values for all genomic 25kb windows for the quartet [*X*]: Fra [*Y*]: Ger [*Z*]: Ira [*O*_1_]: AFG.

**Figure 1–Figure supplement 3.**
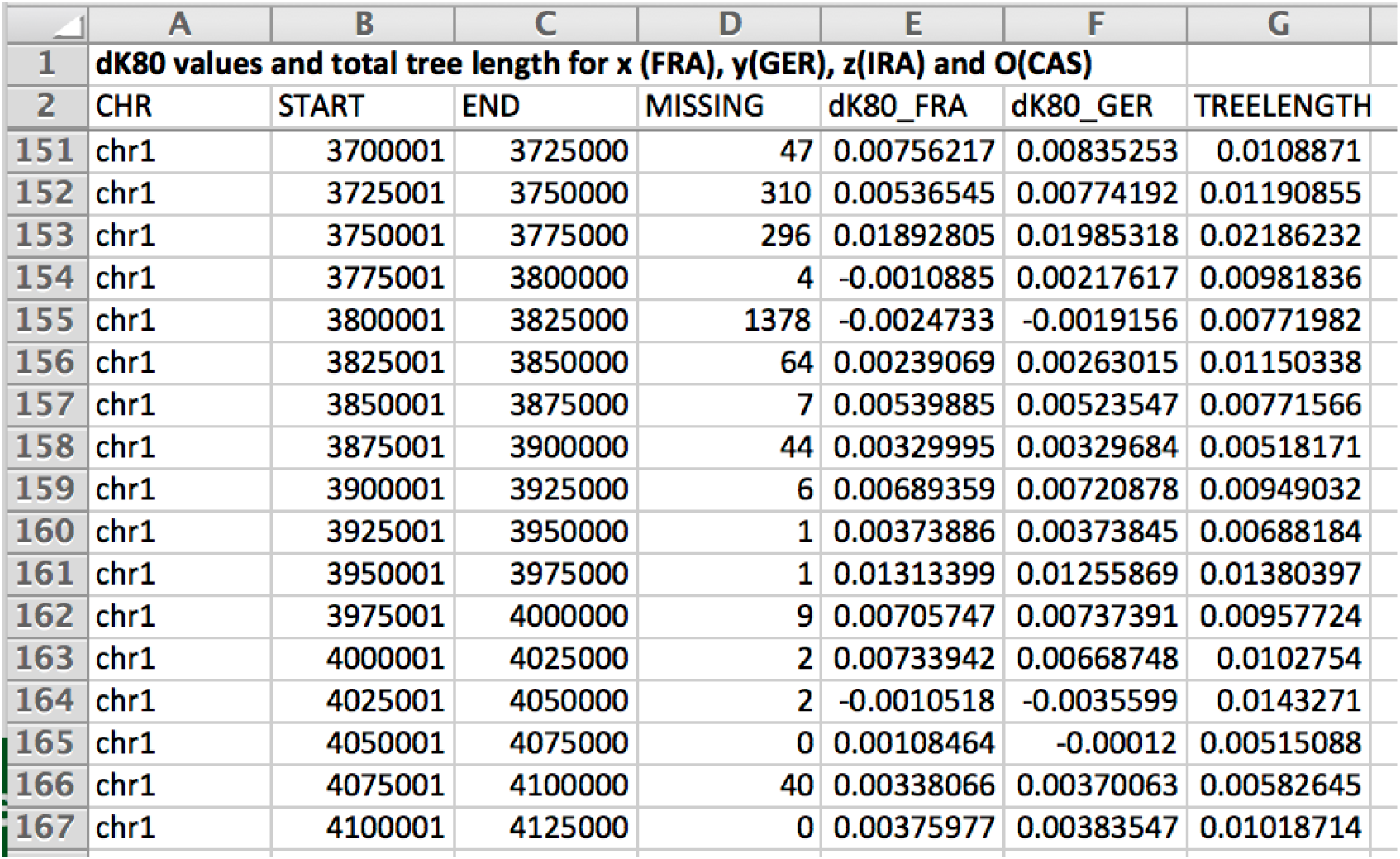
Preview of supplementary Table 2. dK80 values for all genomic 25kb windows for the quartet [*X*]: Fra [*Y*]: Ger [*Z*]: Ira [*O*_1_]: CAS.

**Figure 1–Figure supplement 4.**
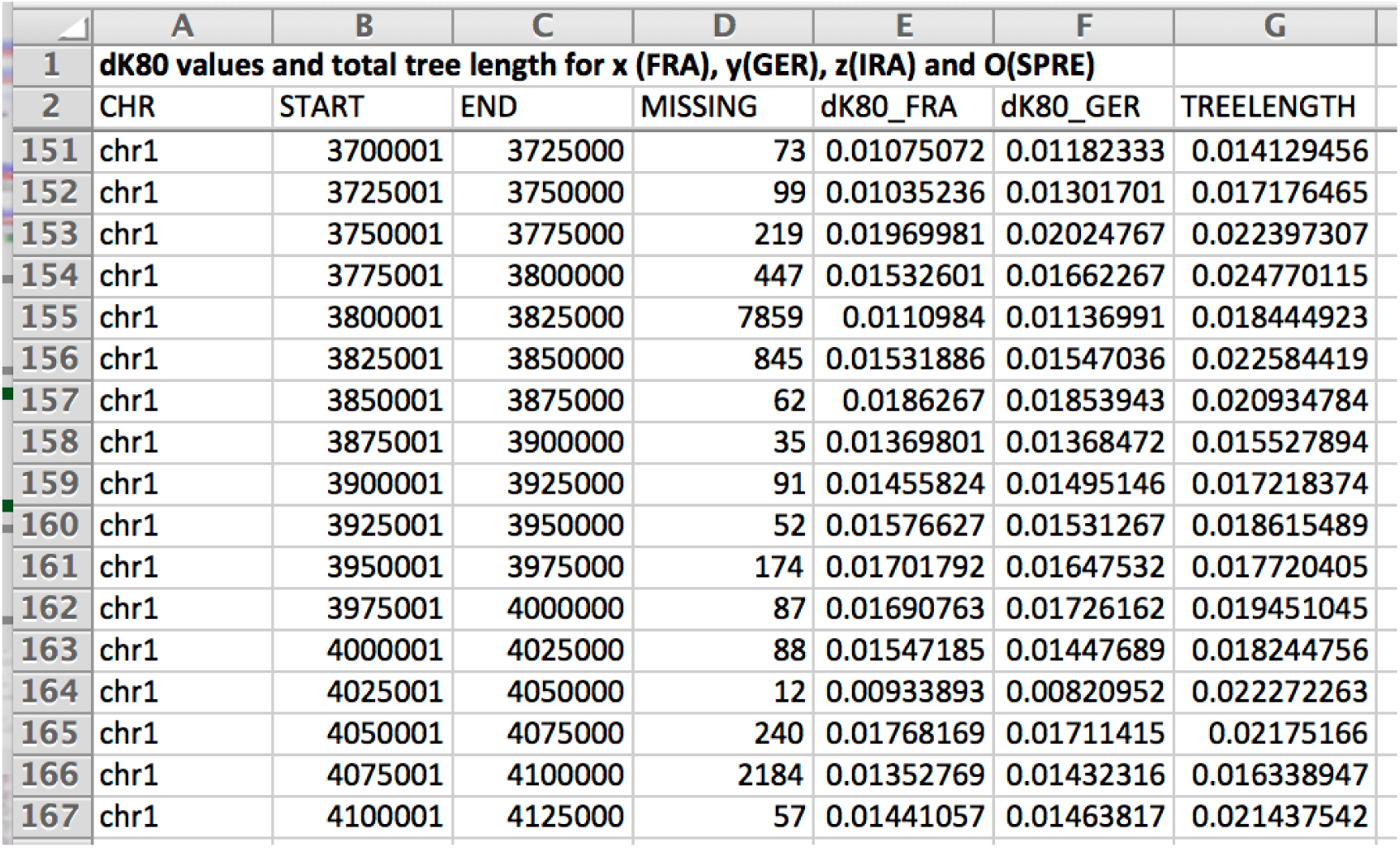
Preview of supplementary Table 3. dK80 values for all genomic 25kb windows for the quartet [*X*]: Fra [*Y*]: Ger [*Z*]: Ira [*O*_1_]: SPRE.

**Figure 1–Figure supplement 5.**
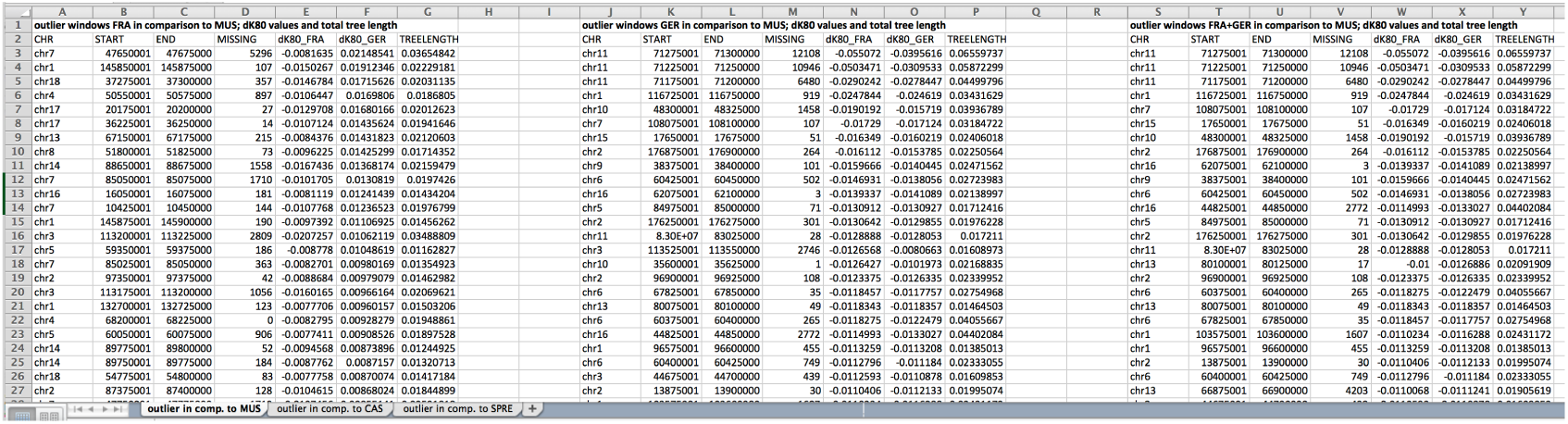
Preview of supplementary Table 4. Lists of outlier windows for the three quartet comparisons.

**Figure 1–Figure supplement 6.**
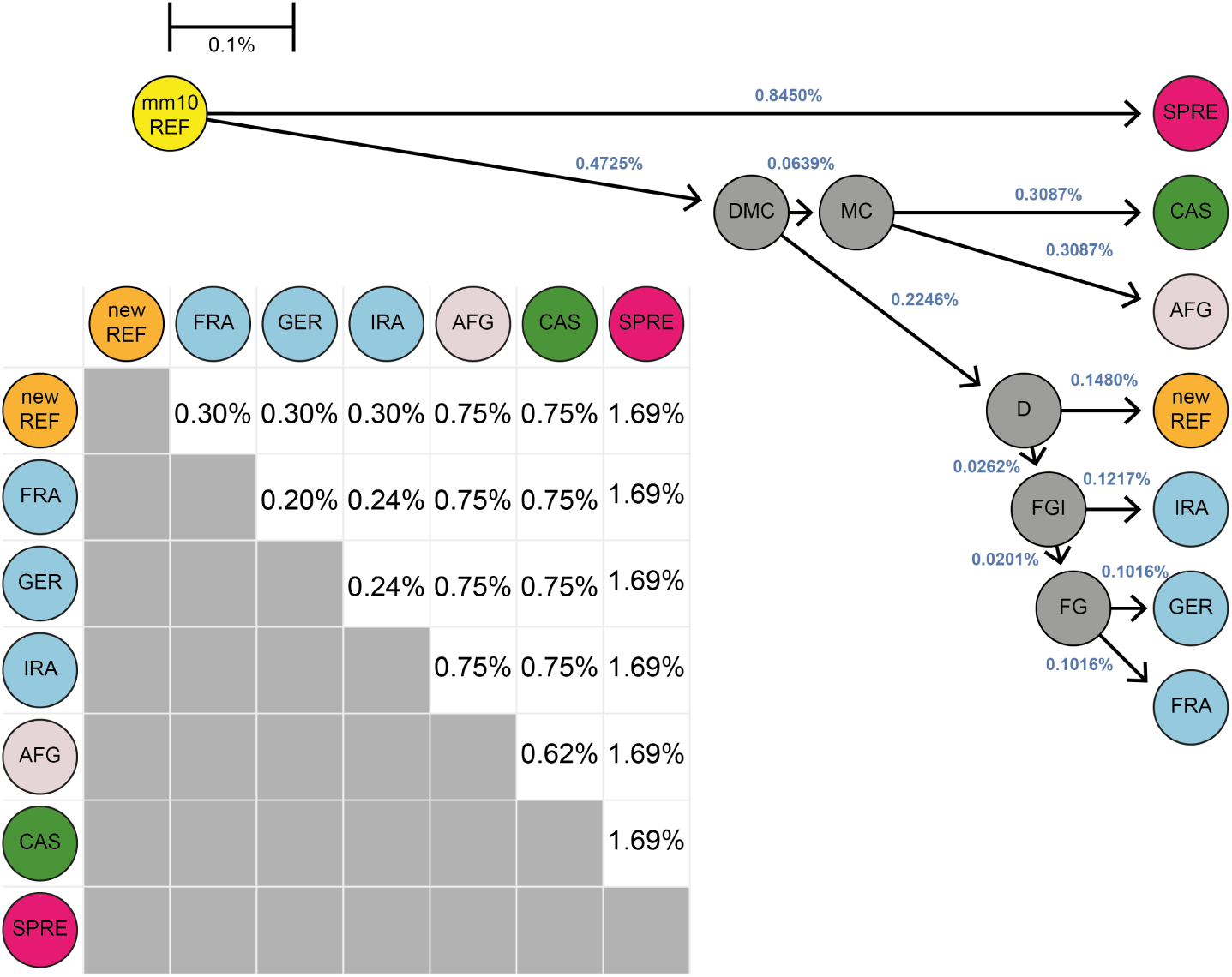
Scheme of the simulation approach. Taking the mm10 reference sequence (yellow) as a start point, genomes were constructed in a phylogenetic context mimicking the real data including the construction of a ‘new’ reference (orange). Nucleotides were randomly altered given a percentage divergence value including ancestral states (grey). The resulting distances represents the phylogenetic context obtained as described in the method section.

**Figure 4–Figure supplement 1.**
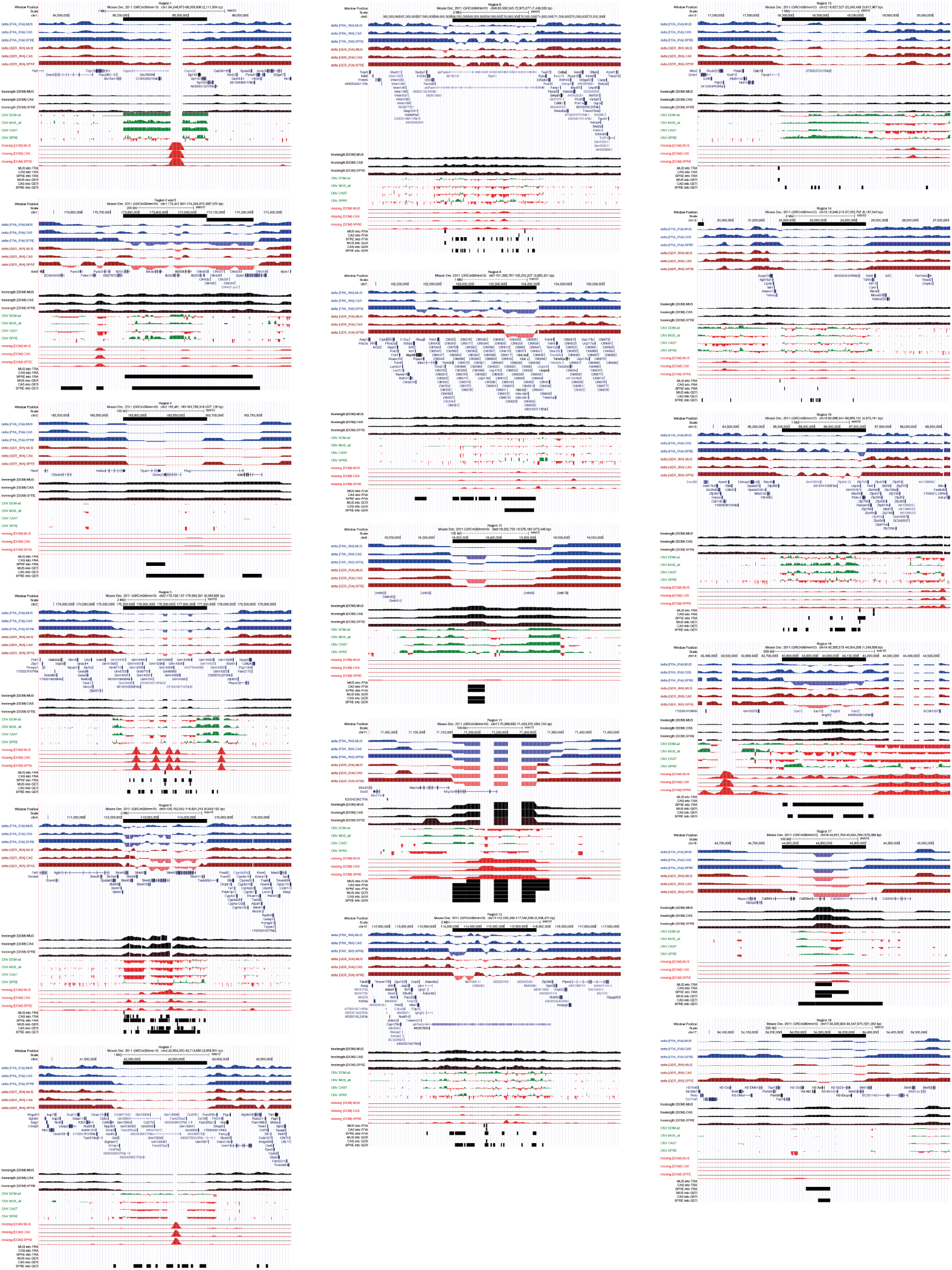
Collection of screen shots of browser windows for all mutual introgression regions listed in Table 2. For full track description see legend of Figure 2.

**Figure 4–Figure supplement 2.**
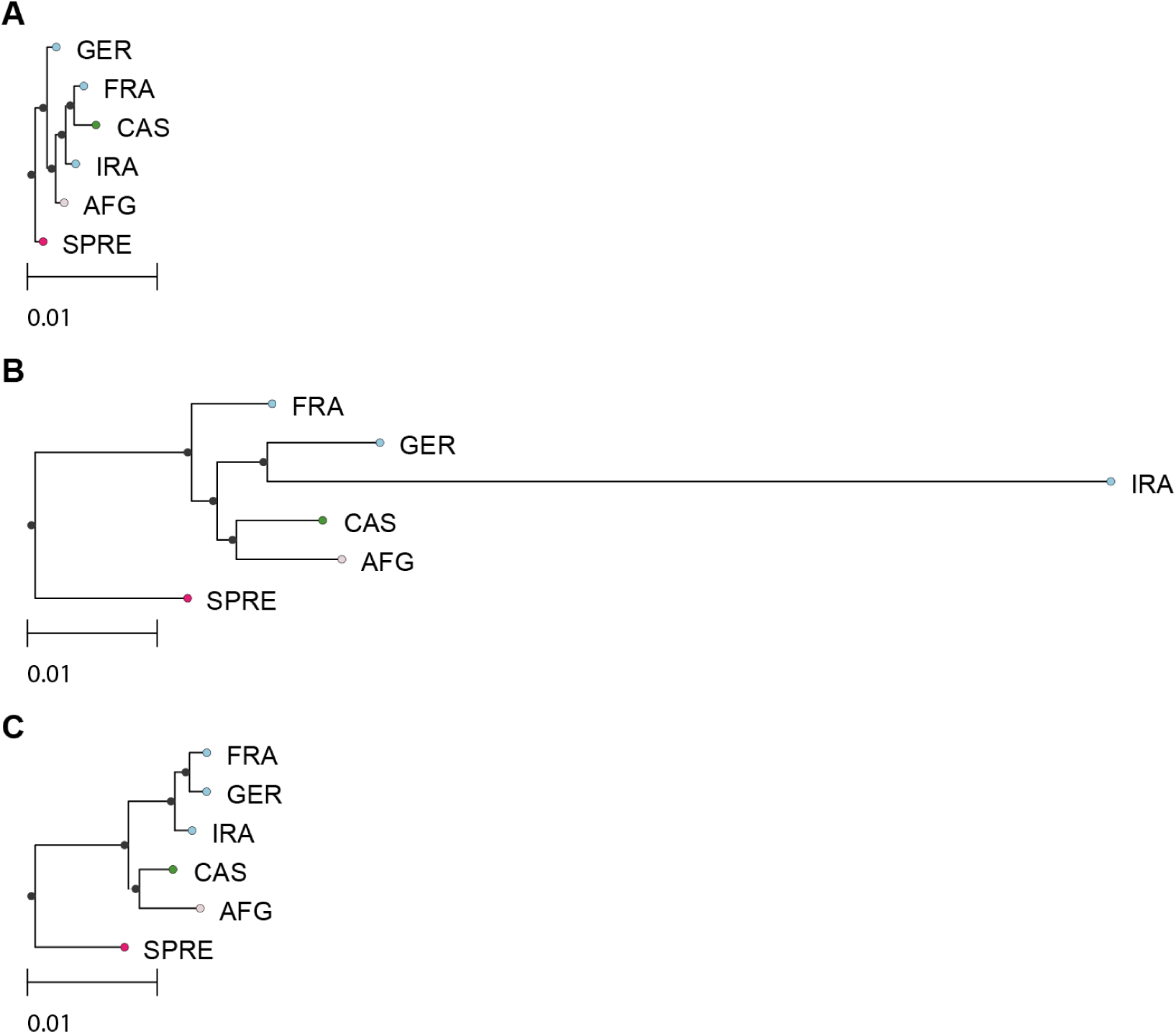
Neighbor-joining (NJ) tree comparisons with consensus sequences in extreme introgression regions. (A) Region 11 from Table 2 (tree constructed for chr11:71,150,213-71,301,792) including *Nlrp1b.* (B) Region 7 from Table 2 (tree constructed for chr4:41,807,517-42,760,683) including a chemokine ligand cluster. (C) Standard tree structure represented by a tree of whole chr19. NJ (Saitou and Nei, 1987) trees based on pair-wise K80 distances (Kimura, 1980) with the R ape package (Paradis et al., 2004). Prior the NJ tree calculation all masked sites were removed from all included sequences.

**Figure 4–Figure supplement 3.**
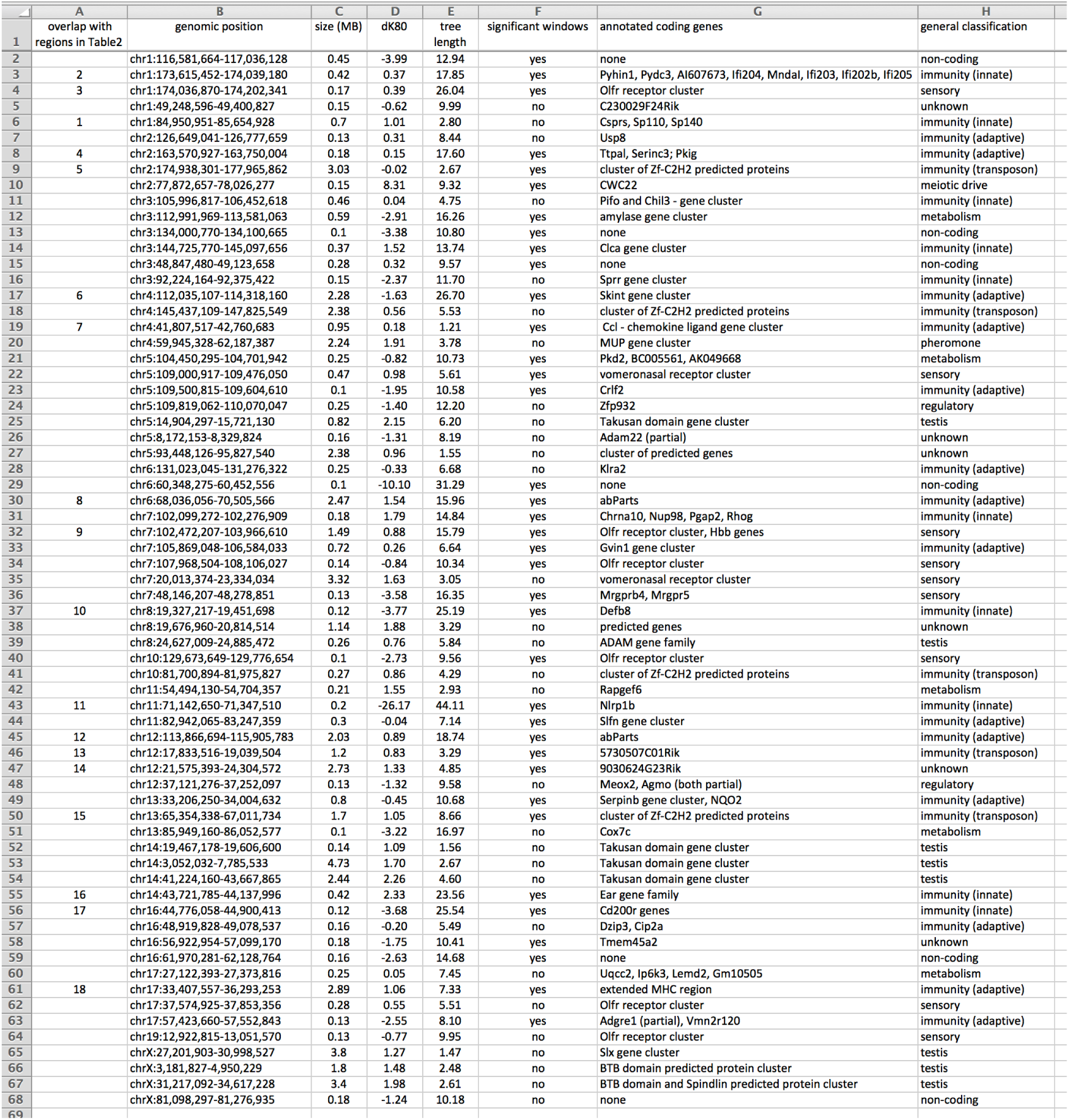
Preview of suppl. Table 5. This table represents an extended version of Table 2 including all identified introgression regions between subspecies and species, as well as the respective genomic locations.

